# Low-cost 3D printed lenses for brightfield and fluorescence microscopy

**DOI:** 10.1101/2023.11.22.568227

**Authors:** Jay Christopher, Liam M. Rooney, Mark Donnachie, Deepak Uttamchandani, Gail McConnell, Ralf Bauer

## Abstract

We present the fabrication and implementation of low-cost optical quality 3D printed lenses, and their application as microscope objectives with different prescriptions. The imaging performance of the 3D printed lenses was benchmarked against commercially available optics including a 20 mm focal length 12.7 mm diameter NBK-7 plano-convex lens used as a low magnification objective, and a separate high magnification objective featuring three 6 mm diameter NBK-7 lenses with different positive and negative focal lengths. We describe the design and manufacturing processes to produce high-quality 3D printed lenses. We tested their surface quality using a stylus profilometer, showing that they conform to that of commercial glass counterpart lenses. The 3D printed lenses were used as microscope objectives in both brightfield and epi-fluorescence imaging of specimens including onion, cyanobacteria, and variegated *Hosta* leaves, demonstrating a sub-cellular resolution performance obtained with low-cost 3D printed optical elements within brightfield and fluorescence microscopy.

## 1. Introduction

Biological imaging research is fundamental in advancing both science and healthcare. To make the resulting developments more widely available both low-cost and open-source microscopy developments can help with circumventing the high economic constraints of conventional commercial biomedical research systems [1]. For some biomedical imaging systems, costs are driven higher due to the requirement for specific optical elements, which can often come at a premium due to their bespoke manufacturing processes and custom design requirements. These factors impose a barrier to entry that constrains biological and diagnostic imaging in low-resource settings. Glass lens manufacturing processes traditionally require grinding and polishing steps, which are time-consuming and costly [2– 4]. Alternatively, injection molded lenses can minimize these costs through mass-scale lens production, however high precision molds must initially be made before high-quality optics can be created, which itself can be both expensive and time-consuming [5–7]. Additionally, when considering the developments of prototype, non-standard and free-form lens geometries employed within imaging research [8–10] in conjunction with the relatively limited customer-base per unique lens specification), the costs in manufacturing each lens or lens mold increases further still. These costs are then passed onto the consumer, slowing, or stopping the participation of biomedical researchers with minimal resources.

Recently, additive manufacturing has demonstrated the potential of 3D printing optical quality free-form lens geometries [11]. Many of the 3D printed optics approaches developed thus far are based on two-photon polymerization (TPP) techniques, such as direct laser writing (DLW), due to their exceptionally high printing resolution and minimal requirements for post-processing [12–14]. However, although TPP can manufacture parts with sub-micron detail, the economic cost per part is significantly higher than other 3D printing methods because expensive pulsed laser sources are necessary for the fabrication process. Additionally, the large spatial footprint and time per print from TPP systems is often a constraint whilst allowing only millimeter-scale parts to be manufactured, albeit with extremely fine detail [12] TPP is an impressive manufacturing technique, it still only caters toward higher-budget research especially for optical quality at the macro-scale.

Stereolithography (SLA) is a low-cost 3D printing technique capable of manufacturing optical components using ultraviolet-excited photopolymerizing resins in a layer-by-layer process [15,16,17]. This stepwise process can require additional post-processing to obtain optical clarity through post-processing approaches like spin-coating or dip-coating with liquid resin, or subtractive methods such as polishing the lens surface with fine grit paper [18,19]. Custom techniques have been developed previously to improve the capabilities of low-cost optical printing, such as iterative learning printing algorithms which adjust grayscale pixel values throughout the print, or image pattern defocusing to smooth the lateral pixel gaps, both of which minimize the occurring staircase effect [20,21], yet still include a coating post-process step. Aside from coating techniques, liquid immersion lens molding can produce lens geometries theoretically invariant of diameter or complexity and without the need for post-processing [22,23]. These low-cost manufacturing methods provide the opportunity to repeatably manufacture optical components with tailored uses within microscopy setups.

Individual lenses or lenslet arrays, manufactured using SLA techniques, have been applied so far to image chrome lithography resolution targets [15,16,24] or by analyzing the shape of a transmitted laser beam [21]. Although these tests are integral for quality benchmarks of in-house lens fabrication, the application and full performance evaluation of 3D printed lenses in biological or biomedical imaging questions has so far been neglected. We therefore present the potential of low-cost LCD desktop 3D printing for optical quality lens manufacturing when used in biological applications. We demonstrate the use of both single and multi-lens microscopy objectives using affordably produced lenses of varying geometries across both brightfield and fluorescence imaging. The imaging performance of each lens geometry was evaluated in brightfield illumination using a 1951 USAF target for resolution and contrast evaluation, as well as cyanobacteria, variegated *Hosta Undulata* leaf, and iodine-stained onion cell samples. We also show fluorescence epi-illumination and imaging using the single lens and multi-lens 3D printed microscopy objectives to demonstrate the potential of 3D printed lenses in biological fluorescence imaging.

## 2. Materials and Methods

### 2.1 3D printing and post-processing protocol

3D printed lens geometries were created in a mechanical CAD software (Autodesk Inventor Professional 2023), with identical geometry to off-the-shelf components for cross-comparison. The design files were translated to printer-readable file formats using a free slicer (Chitubox basic) and printed using an Elegoo Mars 2 consumer grade 3D printer with 10 µm layer step size and lateral pixel size of 50 µm. Clear Resin (RS-F2-GPCL-04, Formlabs) was used as material, with print settings optimized for the material choice and resin material properties having been described previously [25].

The fabrication process to obtain optical quality non-planar components is shown in Fig. 1 A-D. The completed print was cleaned with 100% isopropyl alcohol (IPA) and blow-dried with compressed air before the curved side was spin-coated (Ossila L2001A3-E463-UK spin coater) with a secondary clear resin (UV resin crystal clear, Vida Rosa) with faster cure times. Coating parameters such as spin speed and spin time are intrinsically related to the printed dimensions, such as the surface area and radius of curvature, and were therefore independently optimized for each lens (Table 1). UV curing for 12 minutes was performed using 405 nm and 385 nm LED excitation (Elegoo Mercury Plus 2-in-1 wash and cure station). After curing the curved surface, the lens was again washed with IPA and blow-dried with compressed air.

**Table 1.** Spin-coating characteristics for each 3D printed version of the Thorlabs lenses used in imaging.

| Lens Name | LA1074 | LA1222 | LA1116 | LC2969 |
| --- | --- | --- | --- | --- |
| Coating Speed (RPM) | 1800 | 2800 | 2600 | 4000 |
| Coating Time (s) | 10 | 15 | 14 | 10 |
| Coating Amount (mg) | 120 | 4 | 4 | 4 |

**Fig. 1.**
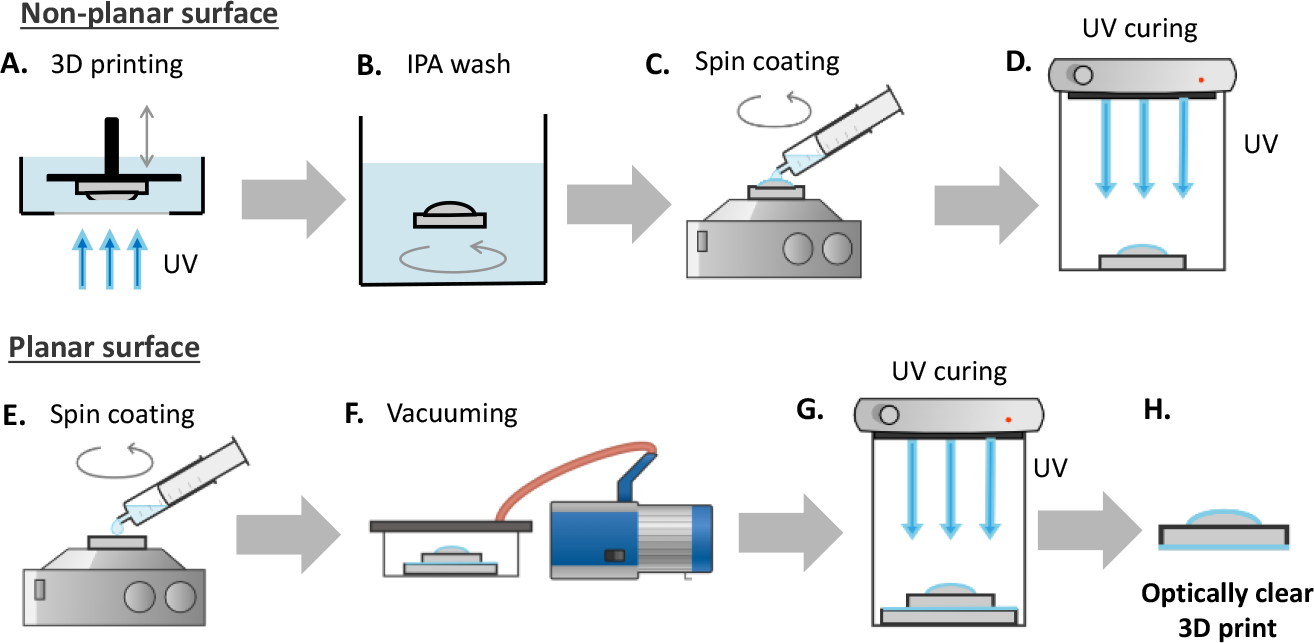
Schematic of 3D printed optics manufacturing process. (A)-(D) Manufacturing an optically clear non-planar surface; (E)-(G) Manufacturing an optically clear planar surface; (A) 3D printing schematic showing the layer-by-layer technique using incident UV light; (B) Cleaning stage using isopropyl alcohol (IPA) to remove residual resin; (C) spin coating non-planar surface by pipetting resin onto surface prior to spinning; (D) non-planar spin coated surface cured with UV light; (E) resin pipetted onto glass slide for spin coating; (F) resin-coated glass slide and 3D printed planar surface placed in contact and vacuumed together; (G) UV curing of resin coated slide and planar 3D printed surface; (H) schematic of resultant optically clear 3D printed part, shown here as a plano-convex lens.

The planar surface coating (Fig. 1 E-H) is created by spin-coating a glass microscope slide with liquid resin (RS-F2-GPCL-04, Formlabs) at 1400 RPM for 10 seconds, and then placing the cleaned planar surface onto the slide to enable best planar surface quality [15].To remove macroscopic air bubbles formed within the resin between the printed lens and glass slide, the combination was left in a vacuum chamber of –0.9 bars for 30 minutes, or until all bubbles had been removed. The lens-slide combination was then UV cured for 8 minutes using the same Mercury Plus wash and cure station as before, and the finished lens was removed from the glass slide using a freezer spray to leverage differential thermal expansion between the printed plastic and glass slide. Some manual leveraging with a scalpel at the 3D printed lens’ edge was additionally required at times. To manufacture and process the described lenses, the material costs were found to be under $0.18 per lens.

### 2.2 Evaluation of surface quality using a stylus contact approach

The optimal spin speed and spin time for the post-process lens coatings were evaluated based on a parametric sweep using a Tencor Alpha Step IQ Stylus profiler with a 5 µm tip to characterize the surface profile of each lens, and a MATLAB script was written to determine any radius of curvature mismatch between the 3D printed lenses and their commercial counterparts.

### 2.3 3D printed single and multi-lens microscope objective design and evaluation

A simple custom microscope setup with fluorescence epi-illumination and brightfield Köhler illumination was used as a testbed for evaluating the imaging performance of the 3D printed optical elements in microscopy applications (Fig. 2). The testbed consisted of a 30 mm optical cage, a 150 mm achromat tube lens (Thorlabs AC254-150-A) and an industrial CMOS camera (IDS U3-3060CP) to capture the test images. Köhler illumination was created through a separate cage system using a white light LED source (Lumiled Luxeon C Starboard) and multiple lenses and apertures to produce uniform illumination with variable numerical aperture. Epi-fluorescence illumination was provided by a multi-mode laser diode module of 488 nm wavelength (Odicforce OFL-488) which was fibre coupled through a 5 m long multimode fibre (Thorlabs M43L05). The output light from the fibre was focused into the test samples using a commercial f = +125 mm achromatic lens (Thorlabs AC254-125-A) and the 3D printed microscope objective under test. For fluorescence imaging, a dichroic mirror (Chroma ZT405/488/561/640rpcv2) was used with two 500 nm long pass emission filters (Thorlabs FELH0500) placed in the infinity space before the tube lens. The 3D printed objectives were placed to create a telecentric imaging setup.

**Fig. 2.**
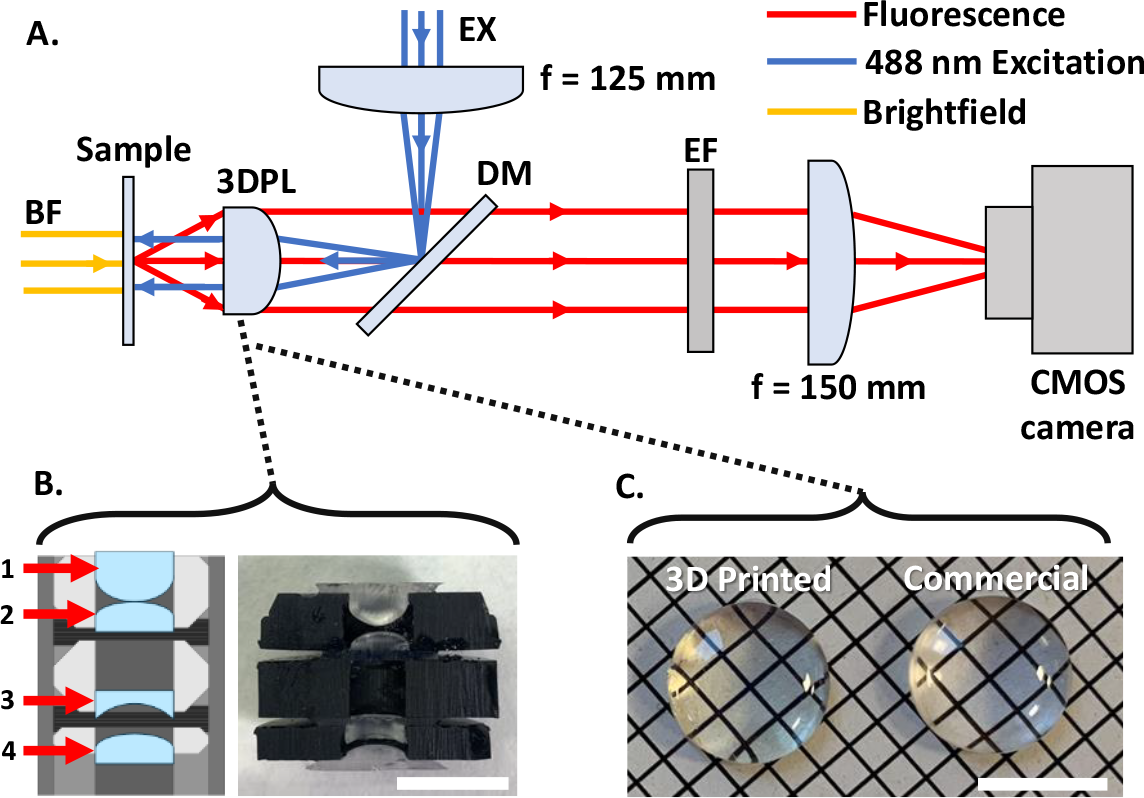
Microscope setup for brightfield and fluorescence image comparison of 3D printed and commercial single and multi-lens objectives. (A) Schematic of the microscope setup using either a group of 4 singlet lenses or a single plano-convex lens as the primary objective. BF –collimated white light source for brightfield illumination; EX – 488 nm collimated multi-mode laser input; 3DPL – 3D Printed objectives under test; DM – dichroic mirror; EF – emission filter. (B) Schematic of 4 singlet lenses and photograph to match schematic. (1) Aspherical f = +6 mm lens; (2) f = +10 mm plano-convex lens; (3) f = -6 mm plano-concave lens; (4) f = +15 mm plano-convex lens. (C) Photograph of 12.7 mm diameter, 3D printed f = +20 mm plano-convex lens and commercial counterpart with identical specifications. Scale bars = 10mm.

The two types of primary microscope objectives using 3D printed elements were designed using the raytracing software Optalix to determine optimal lens placements and configurations. A single plano-convex f = +20mm lens (Thorlabs LA1074) was used as single element objective (see Fig. 2(C) for a comparison of the 3D printed and commercial glass lens). A multi-lens objective geometry was additionally designed based on 6 mm diameter singlet element lenses (see Fig 2B). The objective consisted of an f = +6 mm aspheric front lens (1, Thorlabs APL0606), a f = +10 mm plano-convex secondary lens (2, Thorlabs LA1116), a f = -6 mm plano-concave compensation lens (3, Thorlabs LC2969), and a final f = +15 mm plano-convex lens (4, Thorlabs LA1222). Element spacings were optimized using Optalix, with a theoretical minimum optics limited imaging resolution of 1 µm. The stock lens geometry designs for each lens were taken from the catalogue on the Thorlabs website and used as STL files for printing.

Both the 3D printed and commercial 12.7 mm diameter single element lens objectives were housed in a 30 mm cage plate fixed in a telecentric arrangement in the imaging system, with an 8 mm diameter aperture placed on their convex face to act as an aperture stop to achieve optimal resolution conditions. The commercial and 3D printed 6 mm diameter lenses of the 4-lens objectives were housed in a 12.7 mm lens tube with custom 3D printed spacers, printed at a 50 µm resolution, with chamfered edges and retaining rings which all help to minimize lateral offsets between the lenses and ensures the distance between each lens matched closely the objective design parameters. The minimization of lateral offset errors was essential as the ray trace simulations indicated that displacements >50 µm in lateral directions significantly reduced the achievable image quality. A 2 mm diameter stop aperture was integrated into the 3D printed lens holder design and positioned 1 mm after the final 6 mm lens (Fig. 2B, 4) to avoid overfilling, improving resolution and contrast.

### 2.4 Sample preparation

For brightfield imaging a 1951 USAF resolution target was used (Thorlabs R3L1S4P) along with a sample of variegated *Hosta* (*Hosta Undulata*), a freshwater cyanobacteria sample, and an onion cell sample. For the variegated *Hosta* sample the leaf cuticle was removed with a scalpel. This sample was then rinsed with deionized water to remove any debris that occurred during membrane removal. A small section (5 mm by 5mm) was placed onto a #1.5 coverslip with a small amount of deionized water and sealed with a glass microscope slide. To mount the cyanobacteria, the specimen was washed with deionized water to remove detritus and the cyanobacteria were spread using tweezers on a #1.5 coverslip before being sealed onto a glass microscope slide using nail varnish. The onion cells were prepared by dissecting a 5 mm x 5 mm section of the abaxial epidermis from a brown onion and mounting the membrane (5 mm x 5 mm) onto a #1.5 coverslip with 100 µl of neat Lugol’s iodine solution (62650; Merck, Germany) to produce a stained specimen.

For fluorescence imaging the same *Hosta* and cyanobacteria samples were imaged, making use of the autofluorescence of chlorophyll in the chloroplasts for contrast.

For point spread function evaluation a sample of 500 nm microbeads (Thermo Scientific Fluoro-Max green G500) was used. Five microliters of the aqueous bead solution was placed on a #1.5 coverslip before being sealed with nail varnish on a microscope slide and imaged.

## 3. Results

### 3.1 3D printed lens characteristics

The surface profiles of the 3D printed lenses for the single and 4-lens microscope objective design were measured using the stylus profiler within a cleanroom environment, with the profile of the 20 mm commercial glass polished lens also measured as comparison (Fig. 3A). For the 20 mm lenses the resulting measured radii of curvature are 9.67 mm for the 3D printed lens and 9.94 mm for the commercial glass lens. The shape error for both lenses over a 1.6 mm central diameter is within 200 nm of an ideal spherical shape, with surface roughness below 70 nm RMS for both lenses. For the 6 mm diameter spherical singlet lenses of the 4-lens objective, the measured surface shapes are shown in Fig. 3C and D. The shape error for the 10 mm focal length plano-convex lens (lens 2 in schematic) shows the highest error of almost 4 µm which could be due to the slower spin speeds required for this 6 mm diameter lens in comparison to the plano-concave lens (lens 3 in schematic) which was spun at nearly double the rotational speed. At the same time the error for the plano-concave lens is less than 500 nm. For all three cases the surface roughness is below 150 nm RMS. The measured radii of curvature are for all three cases within 94% of the design spec based on the original Thorlabs design files, with at least three replicates having been measured for each lens type. The flat sides of all lenses have additionally a surface roughness <50 nm [26].

**Fig. 3.**
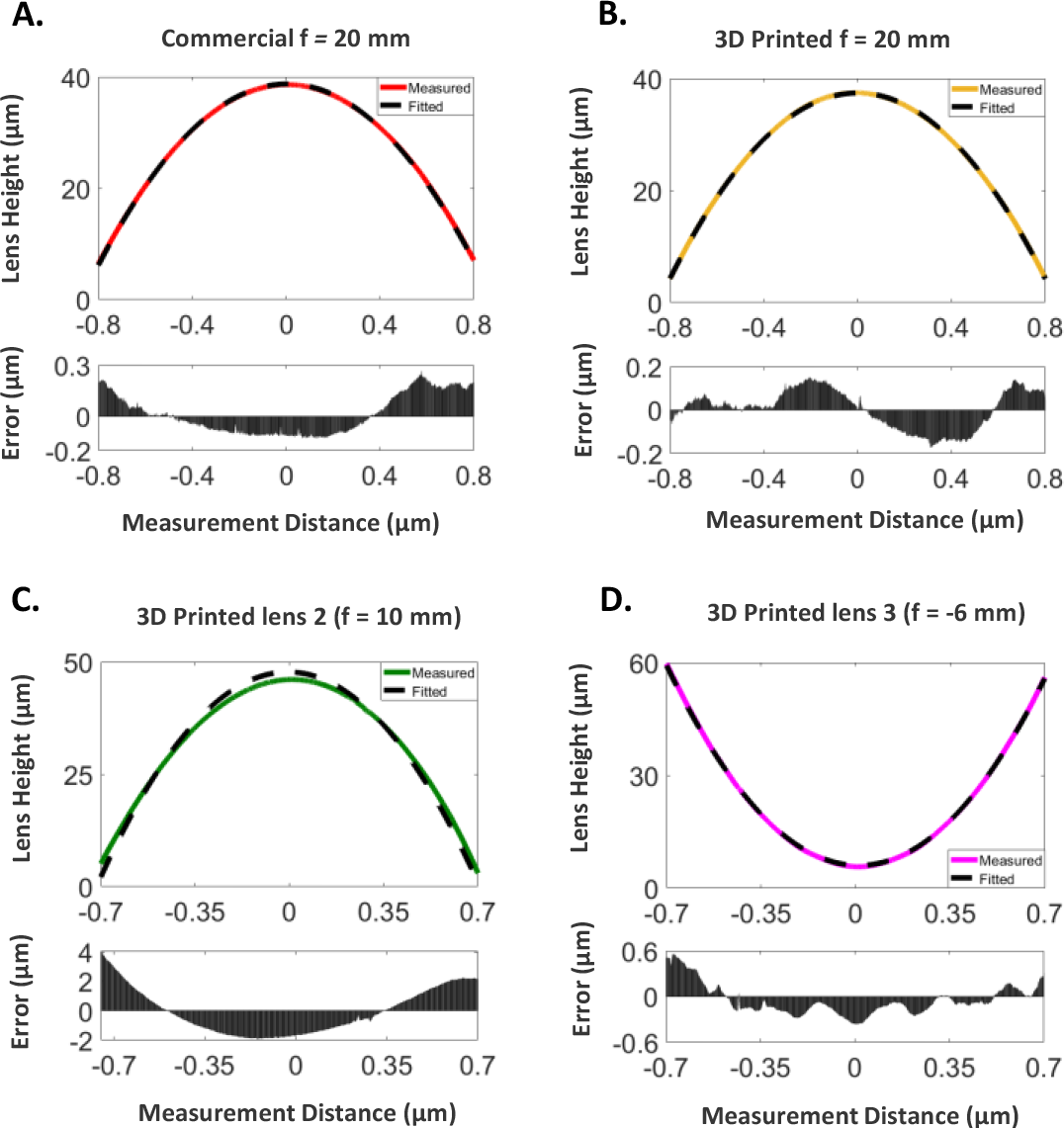
Geometrical characteristics of the 3D printed lenses. (A) Surface profile of 12.7 mm diameter 20 mm focal length commercial lens; (B) Surface profile of 12.7 mm diameter 20 mm focal length 3D printed lens; (C) Surface profile of 6 mm diameter 3D printed lens 2 from the multi-lens schematic; (D) Surface profile of 6 mm diameter 3D Printed lens 3 from the multi-lens schematic.

### 3.1 Brightfield imaging with 3D printed lenses

Using the microscope setup shown in Fig. 2A with the Köhler brightfield transmission illumination and with the single 20 mm focal length lens as the objective leads to an approximately 8X magnification (see Fig.4). The resolution test target shows comparable contrast between the commercial and 3D printed lenses (Fig. 4A and B respectively) over their 1.4 mm by 0.9 mm field of views. For the 3D printed implementation, a small reflection shadow is visible which is attributed to diffraction effects due to voxel patterns within the 3D printed optics. A small degradation in the image quality is visible, with a zoom-in on groups 6 and 7 of the resolution target showing that for the 3D printed lens implementation group 6-6 can be clearly identified, leading to a resolution of around 6 µm, while the commercial lens implementation can resolve down to group 7-1, leading to a resolution of around 4.5 µm. In the digitally zoomed region of interest, the commercial line profile shows slightly higher contrast compared to the 3D printed lens due to slight blurring. Brightfield transmission images of variegated *Hosta*, filamentous cyanobacteria, and onion skin cells stained with iodine (see Fig. 4C-E) show clearly resolvable stomata (see arrow in Fig. 4C) and cell structure for the first case, individual compartments in the cyanobacteria filament, and clearly defined cell walls and nuclei (see arrow in Fig. 4E) in the onion cells for both commercial and 3D printed imaging configurations. The slightly reduced resolution of the 3D printed lens shows extra blurring around the cell walls in the variegated *Hosta*, cyanobacteria and onion compared to the commercial lens images.

**Fig. 4.**
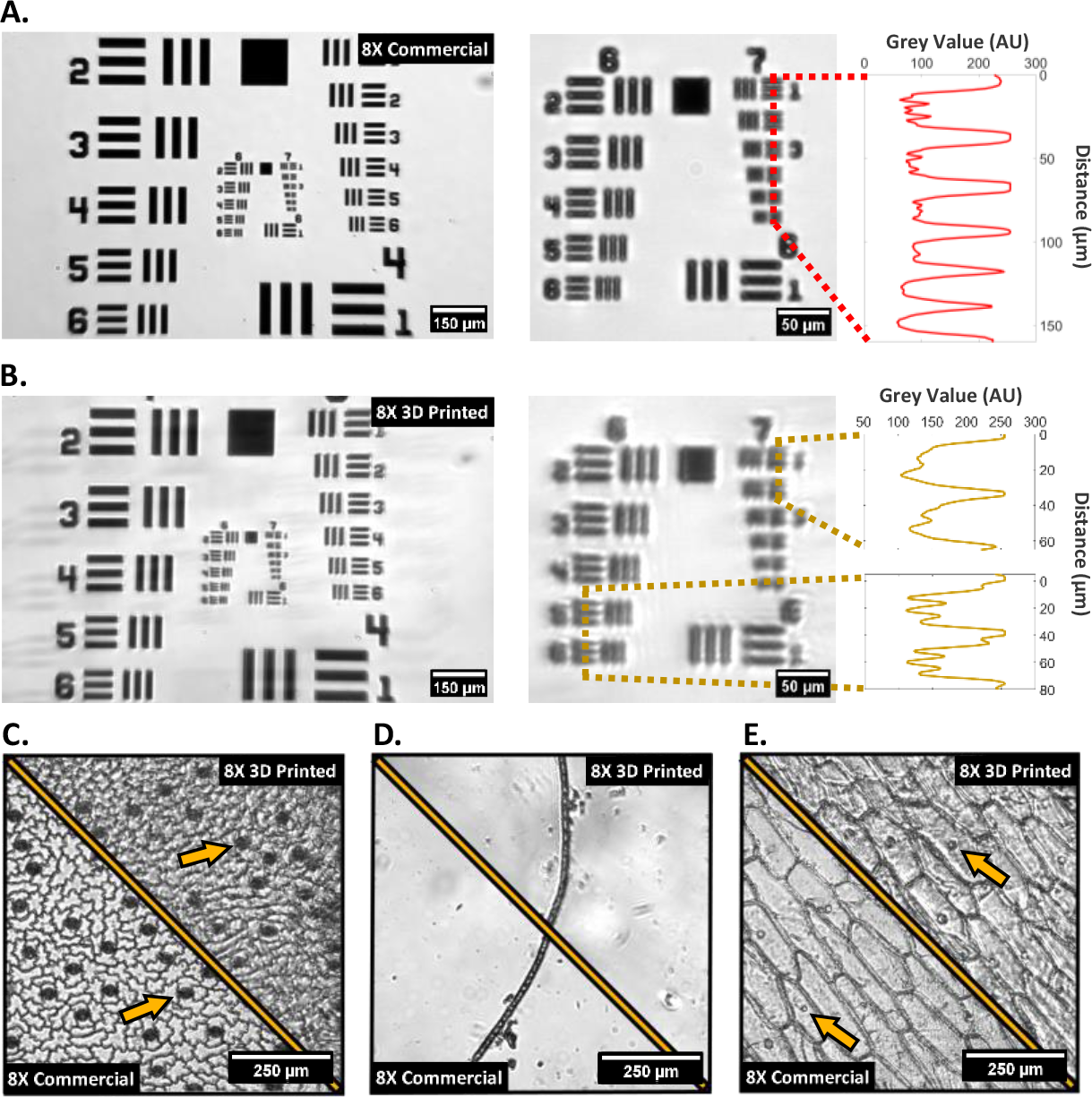
Brightfield imaging using a single f = 20 mm lens as an 8X objective. (A) left -Full FOV 1951 USAF target using commercial 20 mm focal length lens as objective; middle -central cropped ROI from commercial lens image on left; right – line graph taken through group 7 elements 1-6; (B) left -Full FOV 1951 USAF target using 3D printed 20 mm focal length lens as objective; middle -central cropped ROI from 3D printed lens image on left; right - line graph taken through group 6 elements 5-6 and group 7 elements 1-2; (C-E) plant cell images using the commercial and 3D printed 8X configuration of: variegated *Hosta* (C), cyanobacteria filament (D) and iodine stained onion cells (E).

Using the same microscope setup as shown in Fig. 2A, but with the 4-lens objective configuration shown in Fig. 2B instead of the single 20 mm lens, multiple 3D printed 6 mm diameter lenses were used to sequentially replace one or more commercial glass elements to evaluate the impact of using multiple 3D printed elements in succession in a custom microscope objective. Their performance is compared against a completely commercial lens element counterpart using the 1951 USAF target to examine resolution and contrast as shown in Fig. 5A-C. The magnification for the all-commercial implementation is approximately 47X, which is kept consistent when replacing lens 2 (Fig. 2B) with a 3D printed element, however replacing both lens 2 and lens 3 with 3D printed elements increases the magnification to 50X. The shift in magnification is likely a result of sub-millimeter-scale axial displacements of single or multiple lenses in the 3D printed multi-lens setup which, at the high magnification, compounds into the shown differences in magnification. Additionally, diffraction effects are again evident in the 3D printed lens images, shown by faint replicate areas of the target laterally displaced across the FOV. Overall, the use of one or more 3D printed lens elements at the higher magnification provides resolution of the smallest line-pairs on the resolution target, indicating an imaging resolution of <2 µm which allows resolving of sub-cellular details using 3D printed optical elements. With a higher number of 3D printed lenses used in the multi-lens objective, an increased blurring and reduction in resolution is however visible, originating from manufacturing tolerances and potential minor internal scattering within the volume of the 3D printed elements, though these issues were expected from the low-cost manufacturing method.

**Fig. 5.**
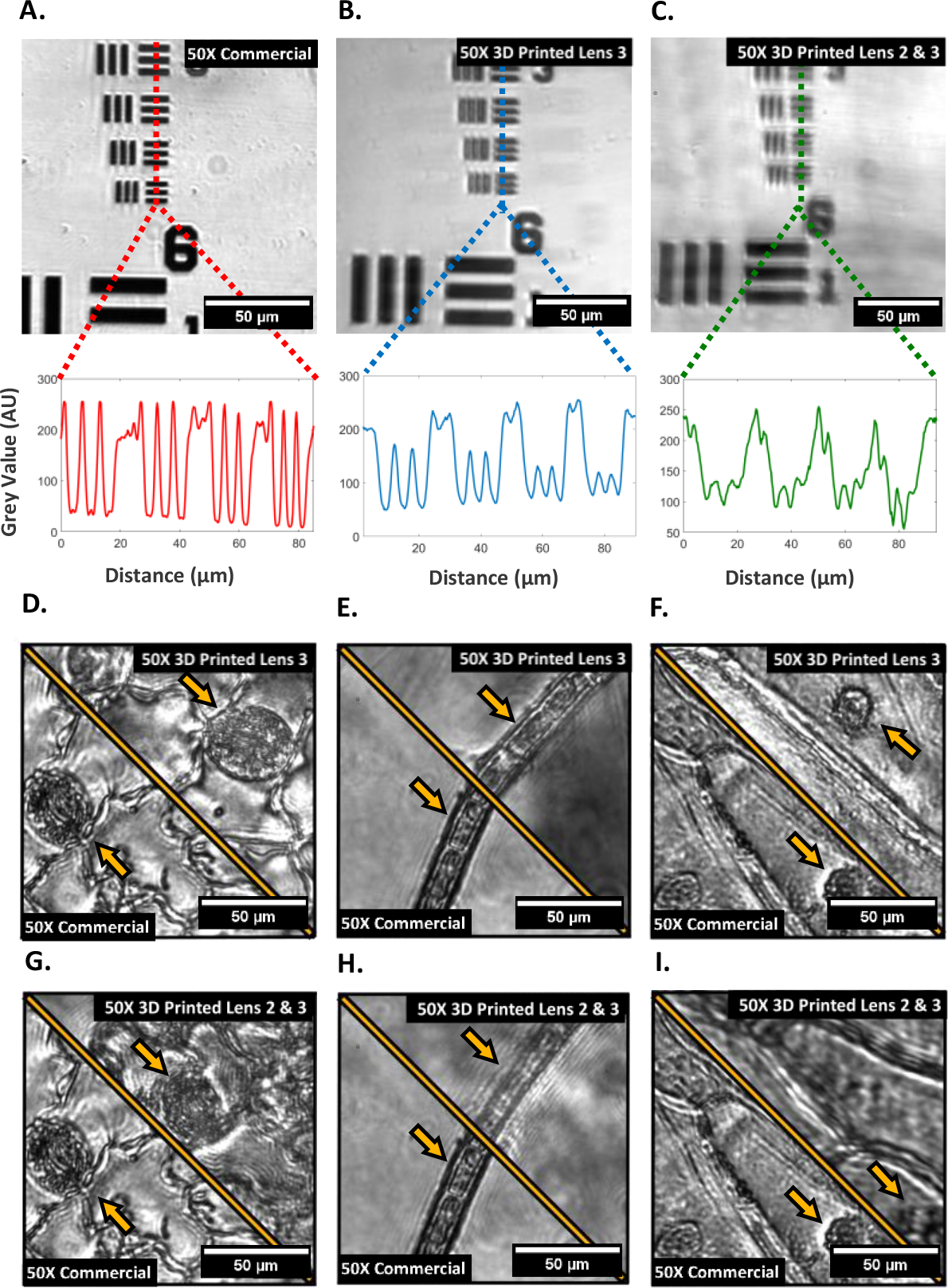
Brightfield images using the 50X multi-lens objective. (A-C) 1951 USAF Group 7 target images with line plot profiles underneath using commercial multi-lens objective configuration. (B) single 3D printed lens (3, plano-concave) objective configuration; (C) two 3D printed lens (3, plano-concave and 2, plano-convex) 50X objective configuration; (D-F) plant cell image comparisons between the commercial 50X configuration and a single 3D printed lens (3, plano-concave) of: variegated *Hosta* stomata (D), cyanobacteria (E) and iodine stained onion (F); (G-I) plant cell image comparisons between the commercial configuration and two 3D printed lenses (3, plano-concave and 2, plano-convex) of: variegated *Hosta* stomata (G), cyanobacteria (H) and iodine stained onion (I).

Fig. 5D-I shows the application of the multi-lens custom objectives with the same biological samples as previously. In these cases, the images of the single and multi-lens 3D printed versions have been scaled to remove the magnification mismatch and allow direct comparison of contrast and image quality. Fig. 5D-F has a direct comparison between an all-commercial implementation and one that replaces a single lens element with a 3D printed version. For all three imaging examples similar contrast can be seen, with minor extra shadowing around the cell membranes due to the mentioned diffraction effects of the 3D printed lens. In all cases, cell nuclei e.g., in the guard cells of the *Hosta* stomata, and membranes (see Fig.5 D-F arrows) are clearly visible and individual compartments of cyanobacteria filaments are distinguishable, including their internal morphology. When adding a second 3D printed lens (Fig. 5G-I) the imaging results show further blurring and potential multiple diffractive effects, which are specifically visible for the single cyanobacteria filament. The contrast and resolution degradation are leading to a reduced visibility of the internal nuclei structure in *Hosta* guard cells and the onion cells, while the main details are still resolvable.

### 3.3 Fluorescence imaging with 3D printed optics

To show for the first time the performance of using 3D printed lens elements in an epi-fluorescence microscope system, images were captured using a 488 nm multi-mode laser excitation with laser illumination and fluorescence collection through the 3D printed elements. A laser excitation power of 500 µW was measured at the sample for the 8X magnification 20 mm focal length commercial and single 3D printed lens objective, and a maximum excitation power of 5 mW for the approximately 50X magnification objectives with multiple 3D printed lens elements. The increased power was necessary to compensate for consistency in the signal to noise ratio between the 20 mm lens fluorescence and the multi-lens fluorescence. Each of the fluorescence images were captured without camera binning at 100 ms exposure time. Using 500 nm sub-resolution fluorescence microbeads, a point spread function (PSF) comparison of the 8X commercial and 3D printed objective configuration is shown in Fig. 6A, together with the PSFs for three multi-lens 50X objective configurations with varying numbers of 3D printed elements in Fig.6B. Significantly degraded PSFs can be seen using the 3D printed lenses as a result of aberrations and scattering effects. The measured fluorescence resolution concurred with Optalix simulations of the PSFs, with the resolution being 3.0 µm and 5.5 µm for the commercial and 3D printed 20 mm focal length objectives, and 0.93 µm, 1.1 µm and 1.4 µm for the commercial, single 3D printed and dual 3D printed multi-lens objectives. To show the biologically relevant application of the two different magnification objectives, samples of variegated *Hosta* and cyanobacteria filament were imaged using the autofluorescence of chlorophyll as the contrast agent (see Fig. 6C-H). Comparing the commercial lens and 3D printed lens for the 8X configuration (Fig. 6C and F) shows good image contrast and the *Hosta* stomata remain easily resolved as can be seen from their characteristic shape and distinct chloroplasts. Additionally, the cyanobacteria has individual cells containing chloroplasts easily discernable from one another in both the commercial and 3D printed cases, though some blurring is observed in the 3D printed version along the edges of the cyanobacteria filament. For the 50X multi-lens objective both the commercial version and the 3D printed version with a single printed lens allow the entire *Hosta* stomata to be resolved (Fig. 6D), with evident sub-cellular detail in the guard cells obtained compared to using the 8X configuration. Though the 3D printed lenses and their commercial counterparts are comparable in resolution, higher blurring is observed through the printed lenses as a result of aberrations and scattering. Similarly, for the cyanobacteria sample (Fig.6G) the commercial and 3D printed lens containing 50X objectives are comparably close in resolution and contrast, though blurring is more prominent in the 3D printed case due to the previously mentioned aberrations and scattering. When adding a second 3D printed lens element into the custom multi-element objective (Fig. 6E and H), a reduction in resolution is visible, specifically looking at the stomata complex in the *Hosta* sample. For the cyanobacteria filament the individual compartments are still clearly visible but the overall clarity of intracellular details is as expected slightly reduced.

**Fig. 6.**
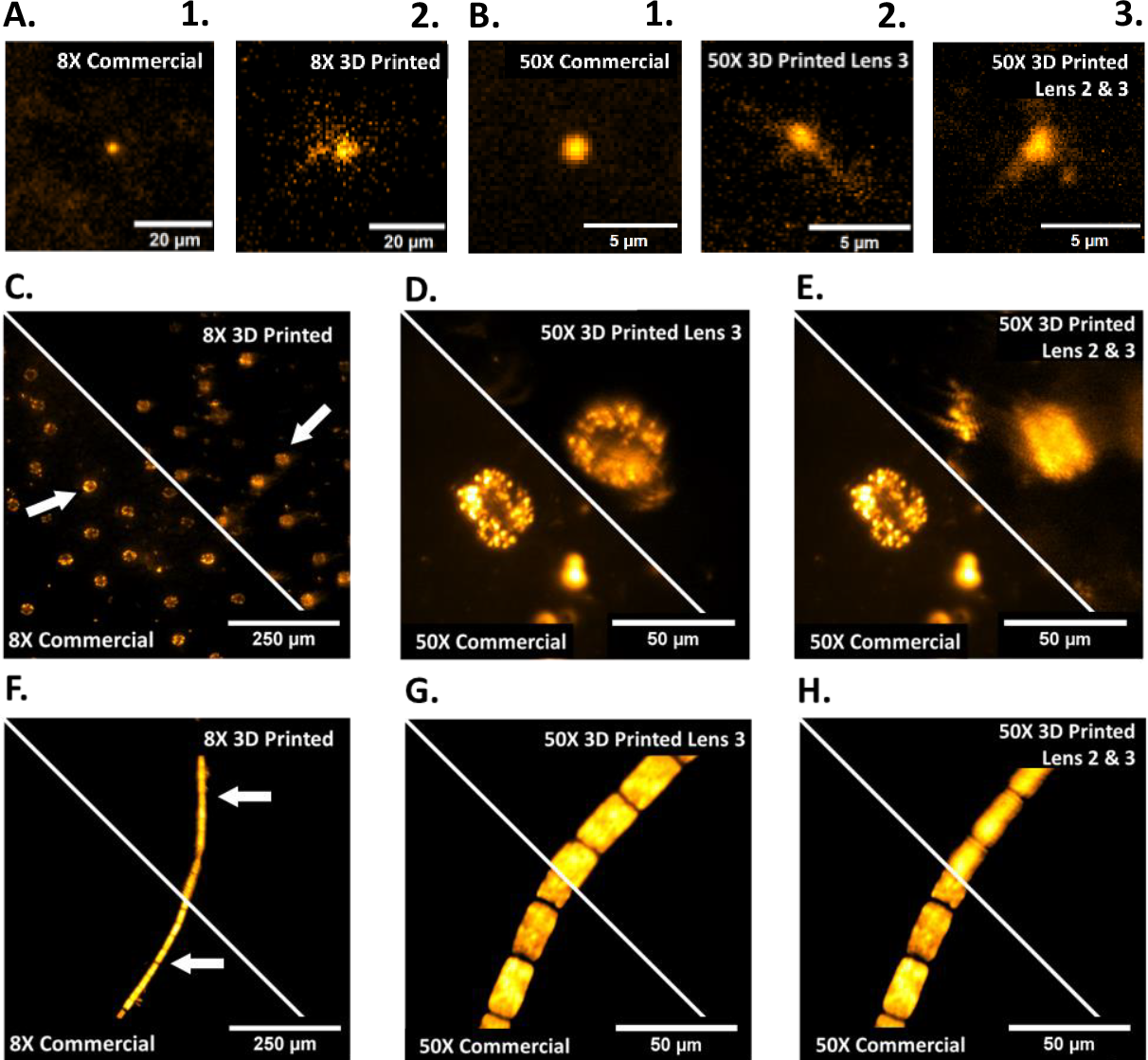
Fluorescence images taken using commercial and 3D printed lens combinations. (A) 500 nm fluorescent beads imaged with the 8X objective configuration. (1) Commercial 8X, (2) 3D printed 8X; (B) 500 nm fluorescent beads imaged with the 50X objective configuration. (1) Commercial 50X, (2) Commercial and 3D printed (3, plano-concave) 50X, (3) Commercial and 3D printed (2, plano-convex & 3, plano-concave) 50X; (C-E) Variegated *Hosta* stomata auto-fluorescence under 488 nm excitation; (C) commercial and 3D printed 8X configurations; (D) commercial and 3D printed 50X (3, plano-concave) configurations; (E) commercial and 3D printed 50X (2, plano-convex & 3, plano-concave) configurations; (F-H) cyanobacteria auto-fluorescence under 488 nm excitation; (F) commercial and 3D printed 8X configurations; (G) commercial and 3D printed 50X (3, plano-concave) configurations; (H) commercial and 3D printed 50X (2, plano-convex & 3, plano-concave) configurations.

## 4. Discussion

As shown in Fig. 4B, faint ‘phantom’ images of the USAF target are observed across the 3D printed lens images, for example visible to the left edge of Fig. 4B where faint replicas of group 4 elements 2-6 can be seen. This has been experimentally determined to be a diffraction effect, likely due to prominent voxel patterning obvious within the uncoated 3D printed lens itself. This generated diffraction grating throughout the print shows that though coating mitigates the staircase effect, the bulk print still influences image quality. The grating pattern exhibited in the uncoated 3D prints may be unique to the used family of printer, as uncoated 3D prints from higher resolution resin printers do not show observable voxel patterns resembling diffraction gratings. The phantom image effects are less obvious in dense samples, in particular Fig. 4C and Fig. 4D, which show more blurred edges likely as a direct product of diffraction though with less discernible laterally displaced ‘phantom’ images. The brightfield images produced of stomata within the *Hosta*, cyanobacteria filament morphology, and onion cell structures using the 8X 3D printed objective are showing similar details to their commercial counterpart albeit with reduced contrast and resolution. For epi-fluorescence applications shown in Fig.6, the 8X commercial PSF seen in Fig.6A shows the expected radial symmetry, while the 3D printed version has scattering and inhomogeneity effects due to the aberrations induced through a spin-coated 3D printed lens and the volume scattering discussed earlier.

When moving to the ∼50X multi-lens objective configuration, it can be seen from Fig. 5 that an individual 3D printed lens at this magnification achieves comparable contrast and resolution to the commercial equivalent, as the finest area of the USAF target are resolved in both cases as well as sub-cellular structural details, for example chloroplasts within the *Hosta* stomata guard cell walls. This is to be expected when considering the quality obtained from the 8X configuration, with aberration effects comparatively concurrent between the two different configurations. Fig. 5 also exemplifies that two 3D printed imaging lenses compound the aberrations which decreases the overall contrast and resolution, though details at micrometer scale are still distinguishable as shown by the USAF target in Fig. 5C. However, Fig. 5G exemplifies that this combination of 3D printed lenses makes identifying individual chloroplasts more challenging in comparison to both the commercial alternative and the single 3D printed lens example in Fig. 5D. When evaluating the ∼50X epi-fluorescence imaging performance, the PSFs in Fig. 6B show similar results to the 8X lenses, where the commercial PSF is radially symmetric as expected compared to the two 3D printed variants which both have significant aberrations. Next to this the PSFs of the 3D printed variants also show significant scattering around the central PSF point, likely originating from the volume grating effect seen in the printed lenses. Additionally, the ∼50X configuration required higher power than the 8X in order to obtain a similar signal to noise ratio, despite being a higher numerical aperture, which can be in part due to the increased number of optical elements within the multi-lens setup.

When comparing 3D printed optical element performance for both brightfield and fluorescence imaging, we observe similar artefacts and aberrations in both the USAF target images in brightfield, and the *Hosta* stomata cell images in fluorescence illumination conditions. So far, 3D printed optics literature has shown resolution target imaging and beam shaping results with comparable resolutions and uniformity to the imaging shown in this work. However, shown for the first time is the application of these manufacturing methods to biological specimen imaging with sub-cellular resolution, drawing attention to the potential for blood smear imaging and therefor low-cost diagnoses of blood diseases and field diagnostics. The low-cost manufacturing method provides ample opportunity for lens combinations as well as custom optics geometries and prescriptions. However, with each 3D printed lens added into the collection arm of the microscope, the contrast and resolution of the imaging system depletes as shown in brightfield (Fig.6) and fluorescence imaging (Fig.7).

## 5. Conclusions

We have shown that using low-cost manufacturing methods for 3D printed optical elements used as single and multi-lens microscope objectives allows collection of both brightfield and fluorescence information in transmission and epi-fluorescence illumination configurations, highlighting their promising potential for custom biological imaging applications. The surface profiles of 3D printed optics closely match their commercial equivalents with curvature radii meeting >90% conformity. While using a single 3D printed lens element in a microscope objective has shown comparable resolution and contrast to commercial glass lenses, we have also shown the impact of adding multiple 3D printed lenses into a custom microscope objective, leading to compounding aberrations in both brightfield and fluorescence imaging. Despite these additional aberrations, we have shown the fluorescence imaging performance using a multi-element 3D printed objective that allowed the resolving of inter-cellular compartments in *Hosta* as well as in cyanobacteria filament samples. The achieved sub-cellular resolution using 3D printed optics within the shown biological samples opens a new gateway into low-cost biological imaging applications in healthcare and field diagnostics.

## Supporting information

Supporting content in the form of ray trace simulations for the two objective families are contained in Supplement 1.

## Funding

UK Engineering and Physical Sciences Research Council (grant EP/S032606/1, studentship EP/T517938/1); the UK Royal Academy of Engineering (Engineering for Development Fellowship scheme RF1516/15/8); the Medical Research Council (grant MR/K015583/1); the Biotechnology and Biological Sciences Research Council (grant BB/T011602/1); The Leverhulme Trust.

## Disclosures

The authors declare no competing interests.

## Data availability

Data underlying the results presented in this paper are available in Ref. [27]

## Notes

https://doi.org/10.15129/d73c68e8-2939-4c24-aa6f-085bf87205d1

### Competing Interest Statement

The authors have declared no competing interest.

### Summary of Updates

Occasional grammatical mistakes fixed; font size and figure sizes decreased; included supplemental material as reference; included one additional reference.

## References

1. B. Diederich, C. Müllenbroich, N. Vladimirov et al., “CAD we share? Publishing reproducible microscope hardware,” Nat. Methods 19 (2022).

2. H. Suzuki, S. Hamada, T. Okino et al., “Ultraprecision finishing of micro-aspheric surface by ultrasonic two-axis vibration assisted polishing,” CIRP Ann. Manuf. Technol. 59, 347–350 (2010).

3. E. Brinksmeier, Y. Mutlugünes, F. Klocke et al., “Ultra-precision grinding,” CIRP Ann. Manuf. Technol. 59, 652–671 (2010).

4. C. F. Cheung, L. T. Ho, P. Charlton et al., “Analysis of surface generation in the ultraprecision polishing of freeform surfaces,” Proc. Inst. Mech. Eng. B. J. Eng. Manuf. 224, 59–73 (2010).

5. J. Luo, Y. Guo, and X. Wang, “Rapid fabrication of curved microlens array using the 3D printing mold,” Optik. (Stuttg) 156, 556–563 (2018).

6. S. Varjo, J. Hannuksela, and O. Silven, “Direct imaging with printed microlens arrays,” Proceedings - International Conference on Pattern Recognition 1355–1358 (2012).

7. L. Li and A. Y. Yi, “Development of a 3D artificial compound eye,” Opt. Express 18, 18125 (2010).

8. R. Wu, Z. Feng, Z. Zheng et al., “Design of Freeform Illumination Optics,” Laser Photon Rev. 12, 1–18 (2018).

9. Z. Feng, L. Huang, M. Gong et al., “Beam shaping system design using double freeform optical surfaces,” Opt. Express 21, 14728 (2013).

10. J. Li, S. Thiele, B. C. Quirk et al., “Ultrathin monolithic 3D printed optical coherence tomography endoscopy for preclinical and clinical use,” Light Sci. Appl. 9 (2020).

11. Q. Ge, Z. Li, Z. Wang et al., “Projection micro stereolithography based 3D printing and its applications,” International Journal of Extreme Manufacturing 2 (2020).

12. S. Wu, J. Serbin, and M. Gu, “Two-Photon Polymerisation for Three-Dimensional Micro-Fabrication,” Journal of Photochemistry and Photobiology A: Chemistry 181, 1–11 (2006).

13. S. Thiele, K. Arzenbacher, T. Gissibl et al., “3D-printed eagle eye: Compound microlens system for foveated imaging,” Sci. Adv. 3, 1–7 (2017).

14. K. Weber, D. Werdehausen, P. König et al., “Tailored nanocomposites for 3D printed micro-optics,” Opt. Mater. Express 10, 2345 (2020).

15. G. D. Berglund and T. S. Tkaczyk, “Fabrication of optical components using a consumer-grade lithographic printer,” Opt. Express 27, 30405 (2019).

16. N. Vaidya and O. Solgaard, “3D printed optics with nanometer scale surface roughness,” Microsyst. Nanoeng. 4 (2018).

17. L. M. Rooney, J. Christopher, B. Watson et al., “Printing, Characterising, and Assessing Transparent 3D Printed Lenses for Optical Imaging,” bioRxiv doi: 2023.11.22.568002 (2023).

18. A. Heinrich, 3D Printing of Optical Components (Springer, 2021), Vol. 233.

19. D. Rodriguez, J. A. Fernández, J. A. Quiroga et al., “Smoothing of 3D Printed Lenses,” US patent 10,086,575B2 (2 October 2018).

20. Y. Shan, A. Krishnakumar, Z. Qin et al., “Reducing lateral stair-stepping defects in liquid crystal display-based vat photopolymerization by defocusing the image pattern,” Addit. Manuf. 52 (2022).

21. Y. Xu, P. Huang, S. To et al., “Low-Cost Volumetric 3D Printing of High-Precision Miniature Lenses in Seconds,” Adv. Opt. Mater. 10 (2022).

22. M. Elgarisi, V. Frumkin, O. Luria et al., “Fabrication of freeform optical components by fluidic shaping,” Optica 8, 1501 (2021).

23. V. Frumkin and M. Bercovici, “Fluidic shaping of optical components,” Flow 1 E2 (2021).

24. G. Shao, R. Hai, and C. Sun, “3D Printing Customized Optical Lens in Minutes,” Adv. Opt. Mater. 8 (2020).

25. M. Reynoso, I. Gauli, and P. Measor, “Refractive index and dispersion of transparent 3D printing photoresins,” Opt, Mater, Express 11, 3392 (2021).

26. J. L. Christopher, P. W. Tinning, and R. Bauer, “3D printing optical components for microscopy using a desktop 3D printer,” Proc. SPIE 12013, MOEMS and Miniaturized Systems XXI, 120130F (2022).

27. J. Christopher, “Data for Low-cost 3D printed lenses for brightfield and fluorescence microscopy,” Univ. of Strath. (2023) 10.15129/d73c68e8-2939-4c24-aa6f-085bf87205d1

