## Supporting content in the form of ray trace simulations for the two objective families are contained in Supplement 1. for "Low-cost 3D printed lenses for brightfield and fluorescence microscopy"

### 1. Ray trace simulations of single and 4-lens imaging system

In this section we present the ray tracing simulations of the two objective configurations used for widefield transmission and fluorescence imaging described in section 2.3 of the main document.

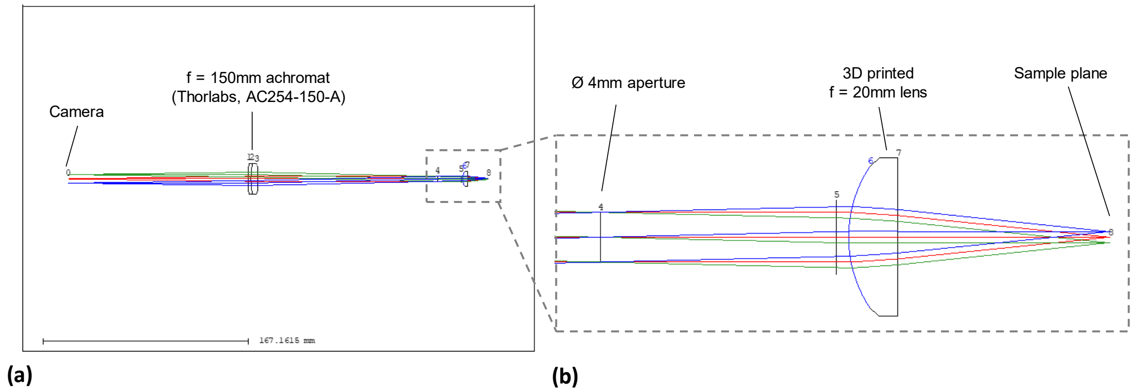


*Fig. S1: Setup and ray propagation of the single 3D printed lens setup. (a) shows the overall setup, with rays originating from the camera chip at the centre and edges of the chip, guided through a f=150mm achromatic doublet lens, a 4mm aperture used as stop aperture and a f=20mm plano-convex lens used as microscope objective. (b) shows a zoom-in of the ray diagrams, focusing on the sample plane and telecentric performance in sample space.*

The configuration using the single f = 20 mm plano-convex lens is shown in Fig. S1, which includes a f = 150 mm achromat used as tube lens for imaging. All rays originate from the camera chip positioned on the left and are traced through the system towards the focal plane of the lens under test. A 4 mm stop aperture is used to balance contrast with achievable imaging resolution, with the overall setup created in a telecentric fashion.

The resulting geometric ray tracing and diffraction analysis is shown in Fig. S2. The diffraction analysis shows a good contrast PSF with small aberrations at the edge of the camera chip and a FWHM of 2.8 µm (Fig. S2(a) and (b)). The three positions shown are the centre of the field of view and edges of the small axis of the IDS U3-3060CP camera chip. The geometrical analysis shows a modulation transfer function (MTF) dropping to zero around 350 Lp/mm (Fig. S2 (c)) and spot diagrams which extend beyond the resolution limited Airy Disk at the centre and edges (see Fig. S2(d)).

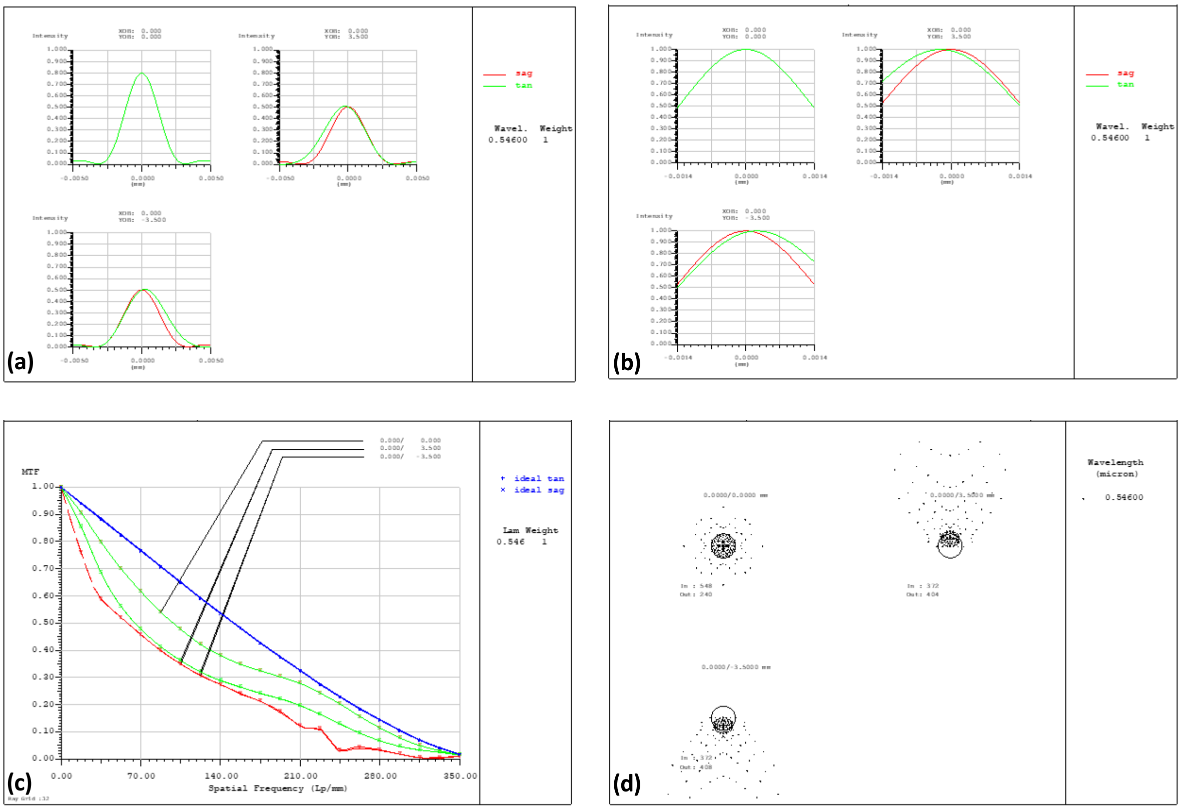

*Fig. S2: PSF, MTF and spot diagram for the single 3D printed lens microscope configuration. (a) and (b) show the point spread function of the system at the sample plane, with (a) showing the contrast degradation at the edge of the field of view and (b) the determination of the FWHM by normalising the PSF graphs. (c) shows the modulation transfer function, highlighting the drop of resolution relative to the diffraction limited case. (d) shows the spot diagram relative to an Airy disk of the system, which highlights that a diffraction limited performance is not achieved over the full field of view.*

The second system demonstrating the use of multiple 3D printed lenses is shown in Fig. S3, including again a f = 150 mm achromat used as tube lens for the system. The objective consists of 4 lenses with 6 mm diameter and an entrance aperture stop with 2 mm diameter. Two plano-convex lenses and a plano-concave lens are 3D printed elements, with a final asphere being an injection moulded plastic lens. For the ray tracing all rays originate again from the camera chip of the used IDS camera and are traced through the system towards the objective stop aperture and the focal plane of the objective under test.

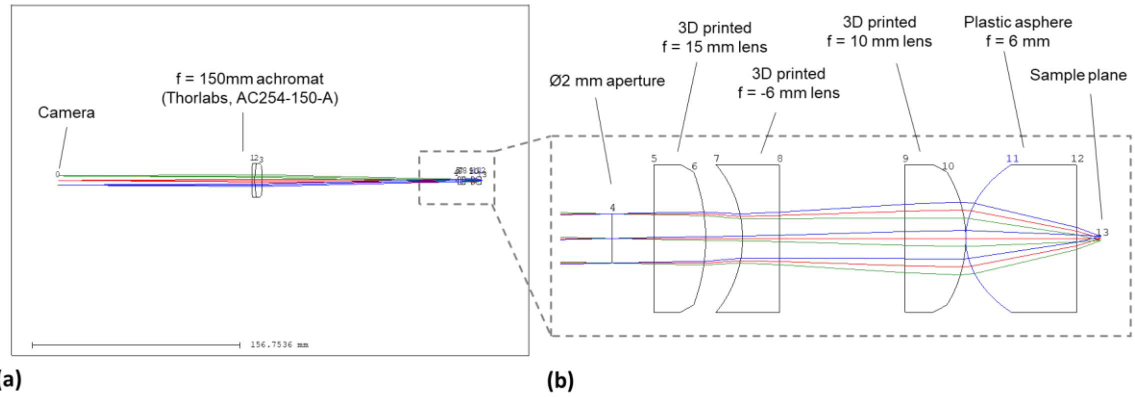


*Fig. S3: Setup and ray propagation of the 4-lens element 3D printed microscope setup. (a) shows the overall setup, with rays originating from the camera chip at the centre and edges of the chip, guided through a f=150mm achromatic doublet lens, a 2mm aperture used as stop aperture and the four lenses used as microscope objective. (b) shows a zoom-in of the ray diagrams, focusing on the sample plane and telecentric performance in sample space.*

The resulting geometric ray tracing and diffraction analysis is shown in Fig. S4. The diffraction analysis shows an excellent contrast PSF with small aberrations at the edge of the camera chip and a FWHM of 1.0 µm (Fig. S2(a) and (b)). The geometrical analysis shows a modulation transfer function (MTF) dropping to zero around 1000 Lp/mm (Fig. S2 (c)) and spot diagrams which at the centre of the field of view are well within the resolution limited Airy Disk and protrude only slightly beyond it at the edges due to the appearing aberrations (see Fig. S2(d)).

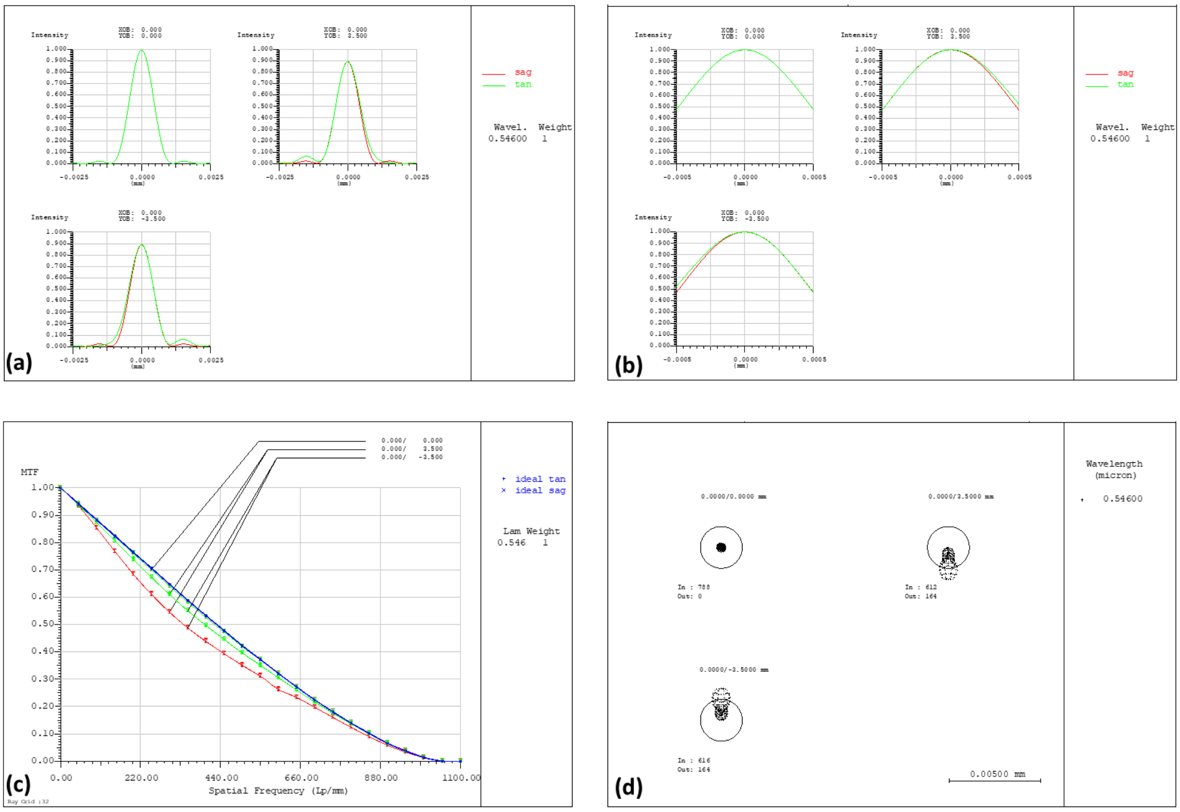


*Fig. S4: PSF, MTF and spot diagram for the 4-lens 3D printed microscope configuration. (a) and (b) show the point spread function of the system at the sample plane, with (a) showing the minor contrast degradation at the edge of the field of view and (b) the determination of the FWHM by normalising the PSF graphs. (c) shows the modulation transfer function, highlighting the small drop of resolution relative to the diffraction limited case. (d) shows the spot diagram relative to an Airy disk of the system, which highlights that a diffraction limited performance is achieved in the centre and over a large area of the field of view.*

Fig. S5 shows the effect of a lateral displacement of the 3D-printed f = 10 mm lens. which has the largest negative impact on the obtainable resolution and contrast. The depicted configuration has a 100 µm lateral offset in y, which shows a clearly reduced contrast in (a), an astigmatic widening of the PSF in (b) and a significant reduction in the MTF graphs relative to the diffraction limit for all three image positions shown in (c). Additionally, the ray tracing shows a significant widening of the ray points in the focal plane, stretching wider than the Airy disk radius for all three image plane points, and highlighting the loss of diffraction limited imaging performance with the introduced offset.

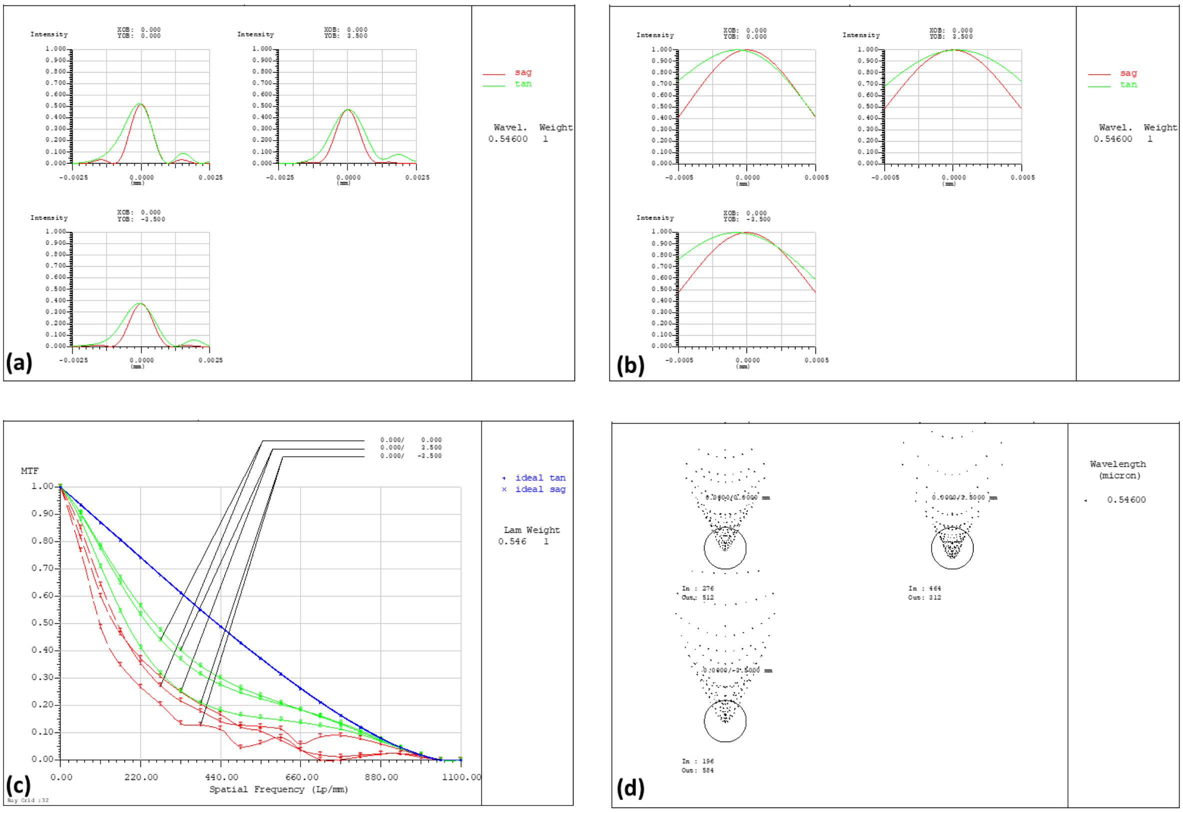

*Fig. S5: PSF, MTF and spot diagram for the 4-lens 3D printed microscope configuration with 100 µm lateral offset of lens 2 (f = 10mm lens). (a) and (b) show the point spread function of the system at the sample plane, with (a) showing the contrast degradation relative to the central placed lens and (b) the determination of the FWHM by normalising the PSF graphs, with significant astigmatism present. (c) shows the modulation transfer function, highlighting the drop of resolution relative to the diffraction limited case. (d) shows the spot diagram relative to an Airy disk of the system, which highlights the degradation of the diffraction limited performance over the field of view.*
